# Assessment of the Genetic Architecture at Early-Stage Drought Tolerance in Wheat Using UAV-Based Multispectral Imaging

**DOI:** 10.1101/2025.11.23.689999

**Authors:** Muhammad Asad Ullah, Fara Muqaddas, Muhammad Kashif Naeem, Atiq ur Rehman, Javaria Tabassum, Shoaib Ur Rehman, Muhammad Sajjad, Zahid Mahmood, Muhammad Arshad Javed, Muhammad Ramzan Khan, Jing Chen

## Abstract

Recent advances in unmanned aerial vehicles (UAVs) and multispectral sensor technologies have transformed high-throughput phenotyping as an efficient alternative to traditional approaches. In this study, we employed UAV-based multispectral imaging to monitor early-stage drought stress in wheat. A panel of 221 historical spring wheat cultivars from Pakistan, representing over a century of breeding history, were evaluated under irrigated and drought conditions. UAV flights were conducted twice at early growth stages to capture multispectral imagery, which was processed in Pix4D mapper for image alignment and orthomosaic generation. Plot segmentation and trait extraction were performed in QGIS. Eight drought-responsive vegetative indices (VIs) were analyzed to assess genotypic variation. Significant to highly significant differences were observed among genotypes, treatments, and their interactions across all VIs. Broad-sense heritability estimates were moderate to high for most traits, with predominantly additive gene action. Vegetative indices such as NDVI, GNDVI, EVI, and SAVI showed strong correlations with each other and effectively detected early canopy stress under drought. Principal component analysis based on 23.897K SNPs indicated a mixed genetic population. Genome-wide association studies identified 115 significant QTNs linked to VIs under both conditions, corresponding to 74 loci, including 26 pleiotropic loci. Of these, 10 pleiotropic loci detected under drought were annotated and six putative candidate genes were identified which showed expression in multiple tissues based on transcriptome data. Two genes i.e., *TraesCS1D01G217600.1* (*HSP70*) and *TraesCS6D01G254000.1* (*ZmMDAR3*), showed differential expression pattern among control and drought treatment. These findings highlight the potential of UAV-based phenotyping for early drought detection and provide candidate genes for improving drought tolerance in wheat. Future research should focus on the functional roles of these genes under drought stress.

## Introduction

A central objective in modern biology is to understand how genotype translates into phenotype. In recent years advances in high-throughput genotyping have made it routine to identify genome-based genetic markers across populations, while phenotyping technologies have not yet advanced at the same pace (Khan et al., 2024). Traditional plant phenotyping methods are often slow, labor-intensive, and may involve destructive sampling. These methods typically capture data at specific time points and contain numerous human-based errors, creating a term that is widely referred to as the “phenotyping bottleneck”. The integration of multidisciplinary approaches, including advanced imaging sensors and robotics, has led to the emergence of high-throughput, large-scale, and non-invasive phenotyping platforms (Zhang and Zhang, 2018, Kumari et al., Yang et al., 2020). The advancement of high-throughput phenotyping (HTP) technologies has significantly accelerated progress in plant phenomics. These systems can capture detailed phenotypic data with high precision and at large scales (Jackson et al., 2023, Chen et al., 2022). The adoption of advanced imaging technologies such as RGB, X-ray, hyperspectral, and multispectral sensors has significantly improved the accuracy of plant phenotyping and contributed to faster progress in molecular breeding (Zhang and Zhang, 2018, Visakh et al., 2024, Sivapragasam et al.). The use of unmanned aerial vehicles (UAVs) equipped with hyperspectral and multispectral cameras has expanded the ability of researchers to monitor large-scale field trials across multiple growth stages within a short time frame (Yang et al., 2020). UAV-derived vegetation indices (VIs) have been effectively used to detect variation among genotypes subjected to biotic and abiotic stress (Khuimphukhieo et al., 2024).

Among the abiotic stresses, drought stress remains a most significant constraint to crop productivity (Ru et al., 2024). Early-stage drought stress in wheat can lead to considerable yield losses, ranging from 18 to 46 percent (Alafari et al., 2024). There is an urgent need to breed drought-resilient varieties that can withstand the environmental challenges using novel genomic approaches like GWAS. Advances in genomics have optimized the efficiency and precision of plant breeding programs via new tools to identify and incorporate drought tolerance QTLs into elite cultivars (Ma and Li, 2024). GWAS have become a key approach for linking genetic variation with complex traits in crops (Uffelmann et al., 2021). The release of the wheat reference genome (Consortium et al., 2018) has further advanced GWAS applications towards precise exploration of genotype–phenotype relationships. This method has successfully identified genes associated with both biotic (Soriano et al., 2021, Wang, 2023) and abiotic stress responses (Gupta et al., 2020), and uncovered candidate loci for important agronomic traits (Pang et al., 2020). In wheat, previously many genomic studies were performed using conventional phenotypic traits at early growth stages (Abid et al., 2016, Dhanda et al., 2004, Yu et al., 2018, Wang et al., 2015). Whereas, HTP for drought evaluation at the early stage under field conditions remains a major challenge. Alongside, robust statistical algorithms and tools are needed to optimize and analyze and comprehend. phenomic data. Over time, VIs derived from spectral bands have been widely used to monitor crop growth and environmental changes (Xue and Su, 2017). Greenness-based indices assess vegetation status, while near-infrared (NIR) bands help estimate leaf water content by minimizing structural effects (Ji et al., 2011). Among these, the NDVI has been the most widely applied for drought monitoring, and long-term VI analysis continues to provide insights into climate variability and drought assessment (Ali et al., 2021). Integration of UAV-based VIs with high-density single nucleotide polymorphism (SNP) data can provide a promising method for conducting advanced quantitative genetic studies and identifying underlying novel loci (Lou et al., 2025, Li et al., 2024). By annotating these loci to genes, they can serve as valuable resources for breeding drought-tolerant varieties. In recent years, image-based phenotyping platforms have been developed to monitor growth and yield-related traits directly from images (Yang et al., 2020). Other studies have shown that detailed image features, such as plant morphology and color, can be used to reliably distinguish between drought-tolerant and drought-sensitive rice varieties (Duan et al., 2018). When combined with GWAS, these phenotyping tools have proven highly effective in identifying the genetic basis of complex traits. For example, dissection of the genetic architecture responsible for stress tolerance in cotton (Li et al., 2017). GWAS found variation in the maize gene *ZmVPP1* to improve seedling survival under drought stress (Wang et al., 2016). Similarly, high-throughput phenotyping platforms employed in rice and found *OsPP15* as a gene contributing to drought tolerance (Guo et al., 2018).

Here, we present an efficient method developed in parallel with existing images processing pipelines. To the best of our knowledge, no previous research has integrated VIs extracted from UAV-based multispectral imaging to investigate the genetic regulation of early-stage drought response in wheat. Therefore, the objectives of this study were to (1) digitally detect drought stress at an early growth stage, (2) identify highly heritable and predictive traits for evaluating early-stage drought response, and (3) uncover genetic factors associated with variation in wheat drought tolerance.

## Materials and Methods

### Field Trial and Experimental Conditions

Field experimentation was conducted during the winter season of 2024–2025 at the experimental area of the National Institute for Genomics and Advanced Biotechnology (NIGAB), NARC, Islamabad, to evaluate a diverse panel of 221 historical wheat genotypes representing over a century of breeding history. The trials were conducted under two contrasting conditions: one with full irrigation and the other with natural rainfed conditions to simulate natural drought.

Wheat genotypes were planted in strip plots with a 1-meter width and a 3-meter length. Sowing was performed using an automated seed drill and seeds of each genotype were pre-weighed at a rate of approximately 25 grams per square meter to maintain a uniform planting density. Each plot consisted of six rows with 25 cm spacing between rows, while a 1-meter buffer was maintained between adjacent strips to minimize interference and allow for accurate phenotypic observations. Standard agronomic practices were followed to ensure optimal crop management.

### Detailed Information Regarding UAV Flights

A Parrot Sequoia drone equipped with RGB, NIR, and red-edge sensors was used for image acquisition. The flight paths were sorted using the DJI GS Pro application on a tablet. Flights were conducted at an altitude of approximately 33 meters, maintaining a ground sampling distance (GSD) of 2.33cm. The frontal and side overlaps were set at 85% and 87%, respectively (Figure 1a). Two flight campaigns were conducted on February 7 and February 28 to target phenotypic response at the early growth stage.

**Figure 1:**
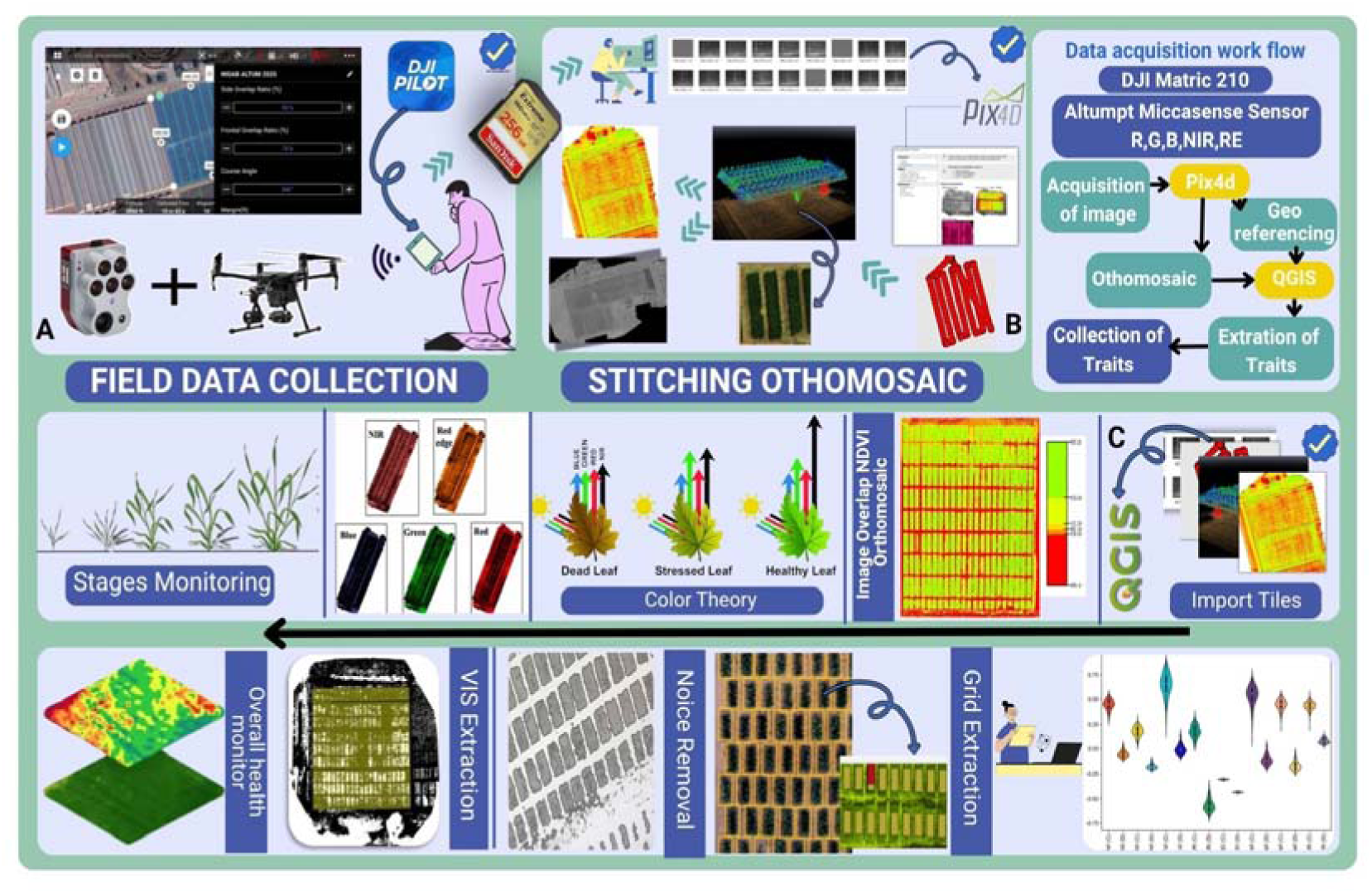
Phenotyping pipeline from image acquisition to trait quantification. A) setting flight map and images acquisition, B) images alignment, 3D dense point cloud and reflectance tiles generation in Pix4D mapper, C) VIs calculation from reflectance tiles, removal of soil noise and masking polygon shapefiles for each plot’s trait quantification.

### Images Processing and Data Extraction

Post-flight, the images from each campaign were aligned and processed using Pix4D (v 4.7.5) software to generate orthomosaics and reflectance maps (Figure1b). During image processing, irradiance correction and radiometric calibration were applied utilizing real-time irradiance measurements from the downward-looking light sensor (DLS) along with reflectance calibration coefficients obtained from the calibrated reflectance panel (CRP). The reflectance maps were then imported into QGIS (v 3.14.16) for VIs extraction (Figure 1c). Eight drought-responsive VIs were computed using QGIS’s raster calculator and then specific thresholds were applied to each index, with values selected manually through visual inspection of the imagery to remove soil noise and weed effect. The NDVI was used as a proxy for chlorophyll content and photosynthetic efficiency. NDREI was applied to assess water status and canopy moisture. The GNDVI, EVI, SAVI, and OSAVI as indicators of vegetation vigor and stress response. The GARI and VARI were used to estimate pigment concentration and detect early signs of oxidative stress. Plot boundaries were delineated manually by overlaying a grid containing rectangular polygons corresponding to each experimental plot. The plot masks were deliberately reduced to 85% of their actual size and precisely aligned within the plot edges to reduce potential edge effects and alignment errors. The zonal statistics tool was used to extract the mean values of each index.

### Phenotypic Analysis of Drought Responsive VIs

Phenotypic trait data were subjected to statistical analysis applying Fisher’s analysis of variance (ANOVA) to determine the significance of differences among genotypes and drought through SPSS software. Variability among traits was further assessed by calculating the coefficient of variation (CV), genotypic variance (σ²g), and phenotypic variance (σ²p). Broad-sense heritability (H²) was estimated using the formula: H² = σ²g/ σ²p. Genetic advance (GA) was computed as: GA = K × σp × H², where K is the selection intensity (assumed as 2.06 for 5% selection pressure) and σp the phenotypic standard deviation.

To visualize the distribution, range, and variability of each trait across time points and treatment conditions, box plots and curve plots were generated using OriginLab. Pearson correlation coefficients were computed using the cor package in R to explore relationships between traits. Principal component analysis (PCA) was performed using the factoextra package in R to classify and group wheat genotypes based on multivariate trait data.

### SNP Profiling and GWAS

A panel of 194 historical wheat cultivars was already genotyped using the 16K genotyping-by-targeted sequencing (GBTS) platform (Majeed et al., 2025). After SNP calling, imputation and QC analyses, a total of 23,897 high-quality SNPs were retained for downstream population genomic and GWAS analyses. GWAS was carried out using a FarmCPU model implemented in the R package rMVP by integrating both genotypic and phenotypic data. PCA and kinship matrix estimation were performed using the “GAPIT” package in R to account for population structure. The model included the first four principal components as fixed covariates to control the population structure and a kinship matrix to account for relatedness among genotypes. Manhattan and quantile-quantile (QQ) plots were generated using the CMplot package in R. SNP-trait associations were considered significant if they exceeded a -logOO(p) threshold of 4.5, highlighted with red dotted line in Manhattan plots.

### Identification of Pleotropic Loci, Favorable Allele Frequency

To identify loci with pleiotropic effects, significant SNPs associated with different traits were compared based on physical position of each chromosome. The pleiotropic loci found under drought condition were further analyzed to detect possible candidate genes. These important and reliable loci were further investigated to identify the favorable allele for each loci using boxplot analysis using OriginLab software. Based on the favorable allele information, we observed the higher accumulation of favorable alleles in different studied cultivars and select the cultivars based on high favorable allele frequency.

### Identification of Putative Candidate Genes and Expression Profiling

To identify the putative candidate genes with respective loci is performed based on proximity-based method. Proximity-based analyses were performed using the Ensemble Plants BioMart tool (http://plants.ensembl.org/index.html) with the bread wheat reference genome (IWGSC RefSeq v2.0). Significant SNPs and its respective position used as reference in BioMart tool and genes located within 2 Mb upstream and downstream of each SNP position were retrieved.

These identified genes were further screened based on gene functional annotation and previous literature information related to drought genes. Further, selected genes were evaluated for transcriptome analysis using expVIP (https://www.wheat-expression.com/) wheat transcriptome database. Genes with differential expression pattern were selected as putative candidate genes. TPM values of these putative candidate genes were normalized and employed for the heatmap in TBtool software. Genes with highest expression level were further observed and validated their role in drought tolerance by qPCR method. Raw expression (CT) values of these genes were estimated using quantitative real-time PCR (qRT-PCR). The *TaActin* housekeeping gene was used to normalized the relative expression data. Relative expression values were evaluated by using the 2^ΔΔ^CT method. Bar-plots were developed in excel using 2^ΔΔ^CT and standard error values.

## Results

### Optimizing Field-Based UAV Multispectral Scans for Assessing Drought Responses in Wheat

UAV-based multispectral imaging proved highly effective for evaluating drought responses in wheat as a non-destructive phenotyping of a diverse panel of historical genotypes. The UAV was equipped with sensors with an average focal length of 1,445.20 pixels (equivalent to 5.40 mm) to capture reflectance data across five spectral bands: NIR, red, green, blue and red-edge. The images were acquired at a fine ground sampling distance (GSD) of 2.33 cm. Dense point clouds with 15 overlapping images were used to construct 2D mosaic maps, with an average reprojection error of 0.21 pixels. A total of 35 GB of multispectral image data was collected during UAV flights over 884 plots, completed within a 1.2-hour total flight time. The raw imagery was processed using machine learning techniques to extract vegetation indices (VIs) from reflectance data for assessing key traits at the early growth stage.

### Dynamic genotypic and treatment effects on phenotypic variation

Differences in significance pattern were observed across vegetation indices in response to genotypic and treatment effects. NDVI showed significant variation among genotypes and highly significant effects of treatments and genotype × treatment interaction. GNDVI, EVI, NDREI, and OSAVI all exhibited highly significant variation due to genotypes, treatments, and their interaction. GARI displayed highly significant variation among genotypes and for the genotype × treatment interaction, while the treatment effect was significant. In contrast, VARI showed highly significant variation among genotypes and their interaction with treatments, but no significant response to treatment.

Vegetation indices derived from multispectral imagery revealed consistent and significant differences between control and drought-treated wheat genotypes across both time points (Figure 2). NDVI and GNDVI values were markedly lower under drought conditions, indicating reduced chlorophyll content and photosynthetic efficiency, with further declines observed from TP1 to TP2. EVI also showed substantial reductions in drought-exposed samples, suggesting a loss of canopy structure and biomass over time, while control plants maintained stable, higher values indicative of healthy vegetation. GARI values sharply decreased under drought, turning negative and signalling severe physiological stress and pigment degradation, in contrast to the positive values maintained by control samples. Similarly, NDREI values were significantly lower in drought-stressed genotypes. SAVI and OSAVI, both indicators of vegetation vigor and greenness, were drastically reduced under drought, with limited variability and values clustering near zero, whereas control plots maintained consistently higher and more variable readings. Overall, the indices demonstrated strong discriminatory power in capturing drought-induced physiological and structural changes in wheat.

**Figure 2:**
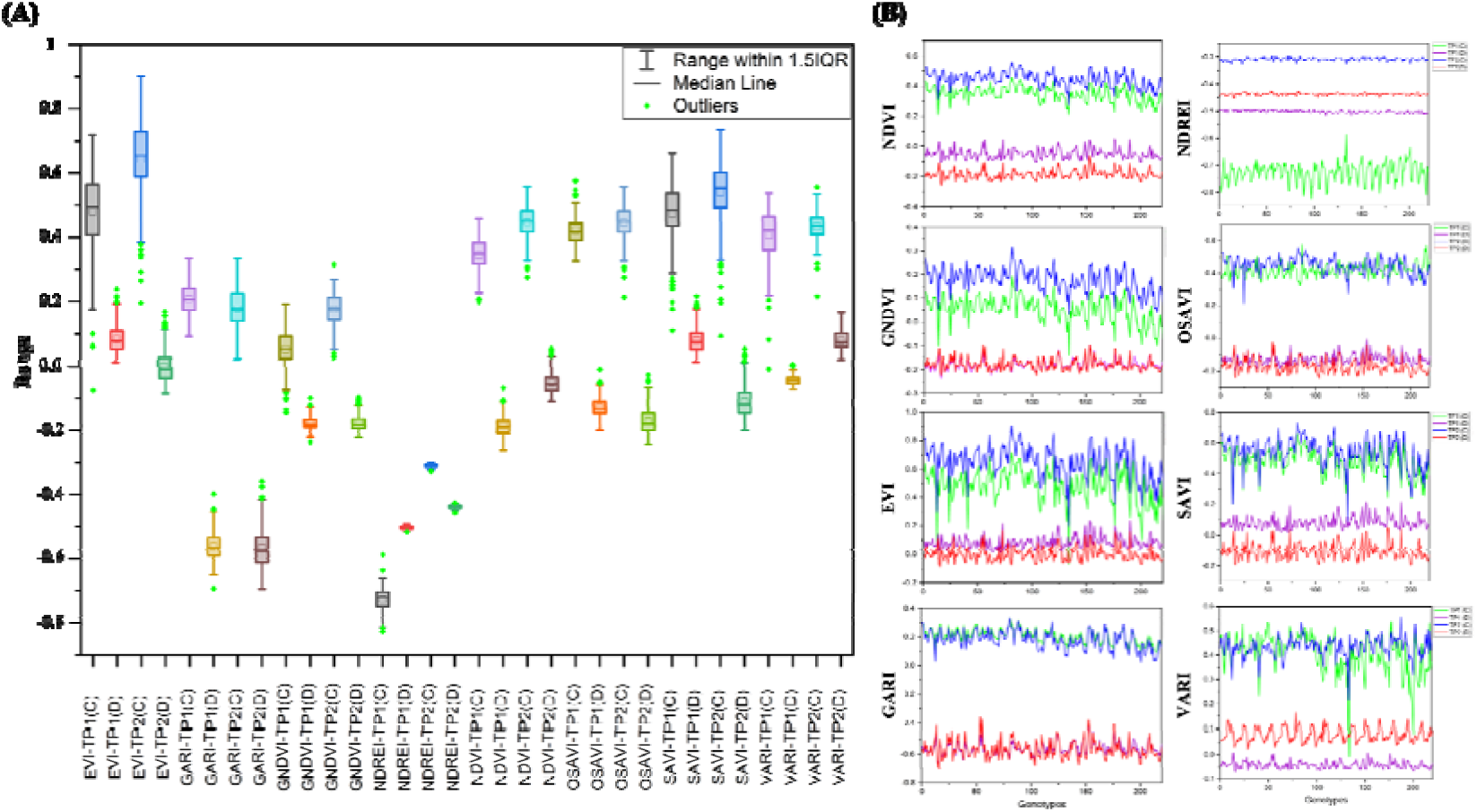
Phenotypic variations for eight vegetative indices estimated across two time points. A) Box plot showing variation of VIs, where C represents irrigated condition and D is for drought condition, B) curve plots showing variation in VIs values for each genotype.

### Genetic Dissection

The analysis of genetic variability revealed that GNDVI exhibited the highest genotypic coefficient of variation (GCV%) at 58%, followed by NDVI (44%) and GARI (41%), indicating substantial genetic diversity and strong potential for selection within these indices (Figure 3a). In contrast, OSAVI and SAVI displayed the lowest GCV values, at 35% and 37% respectively, suggesting more limited genetic variation. The phenotypic coefficient of variation (PCV%) was highest for SAVI (54%), OSAVI (52%), and NDREI (50%), reflecting the influence of both genetic and environmental factors on these traits. NDVI and NDREI showed the lowest environmental coefficient of variation (ECV%) at 8%. In comparison, GARI and OSAVI recorded the highest ECV values (13%).

**Figure 3:**
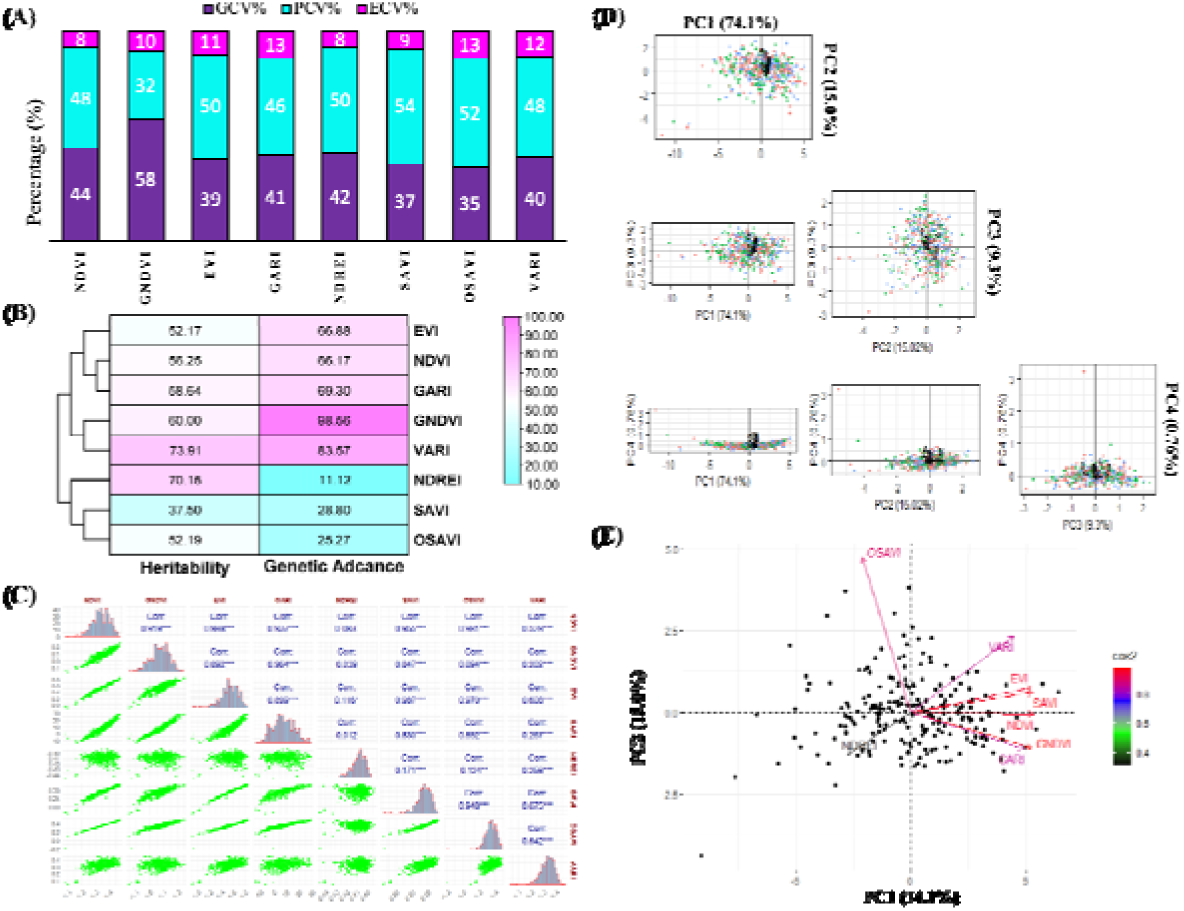
Genetic variability for VIs and classification of diverse genotypes. A) Bar chart for GCV, PCV and ECV of all VIs, B) heatmap of heritability and genetic advance of all VIs, C) association among the eight studied VIs, D) PCA plots for multiple PCs, E) PCA biplot for first two PCs.

Broad-sense heritability estimates ranged from 52.17% to 70.14%, with NDREI demonstrating high heritability, while the remaining indices exhibited moderate levels (Figure 3b). Genetic advance was high for most indices (predominantly controlled by additive genetic effects, indicating the potential for effective improvement through selection), while NDREI showed moderate genetic advance despite its high heritability that might be influenced by non-additive gene effects such as dominance and epistasis.

### Association of Different VIs Among Each Other

Pearson correlation analysis revealed that NDVI showed very strong positive correlations with GNDVI, EVI, GARI, and SAVI, and a strong correlation with VARI (Figure 3c). GNDVI also correlated strongly with EVI, GARI, SAVI, and VARI, but exhibited moderate-to-strong negative correlations with NDREI and OSAVI. EVI was tightly associated with SAVI and strongly linked to GARI and VARI, while moderately negatively correlated with NDREI and weakly with OSAVI. GARI followed a similar pattern, positively linked to SAVI and VARI, but negatively to NDREI and OSAVI. In contrast, NDREI showed moderate negative correlations with most indices except for a weak positive correlation with OSAVI. SAVI was positively correlated with NDVI, GNDVI, EVI, GARI, and VARI, and weakly negatively with OSAVI. OSAVI, unlike other indices, exhibited moderate-to-strong negative correlations with most traits and was uncorrelated with VARI. VARI was strongly aligned with indices linked to vegetation greenness and structure.

### Classification of Genotypes and VIs based-on PCA-biplot

PCA revealed that the first two principal components, PC1 and PC2, explained 74.1% and 15.02% of the total variance, respectively, accounting for a cumulative 89.12% of the overall variability (Figure 4d). NDVI, GNDVI, EVI, SAVI, and GARI clustered closely with long vectors pointing strongly along the positive axis of PC1, indicating that these indices are highly interrelated and contribute substantially to the main axis of variation. NDREI, with a shorter arrow pointing opposite to NDVI and EVI, showed a weaker association with PC1 and a negative correlation with most other indices. The distribution of genotypes across the biplot was concentrated near the origin, indicating that many genotypes exhibited average variation. However, the spread along specific vector directions, such as those of NDVI/EVI or OSAVI/VARI, reflects genetic diversity in how individual genotypes express different VIs.

**Figure 4:**
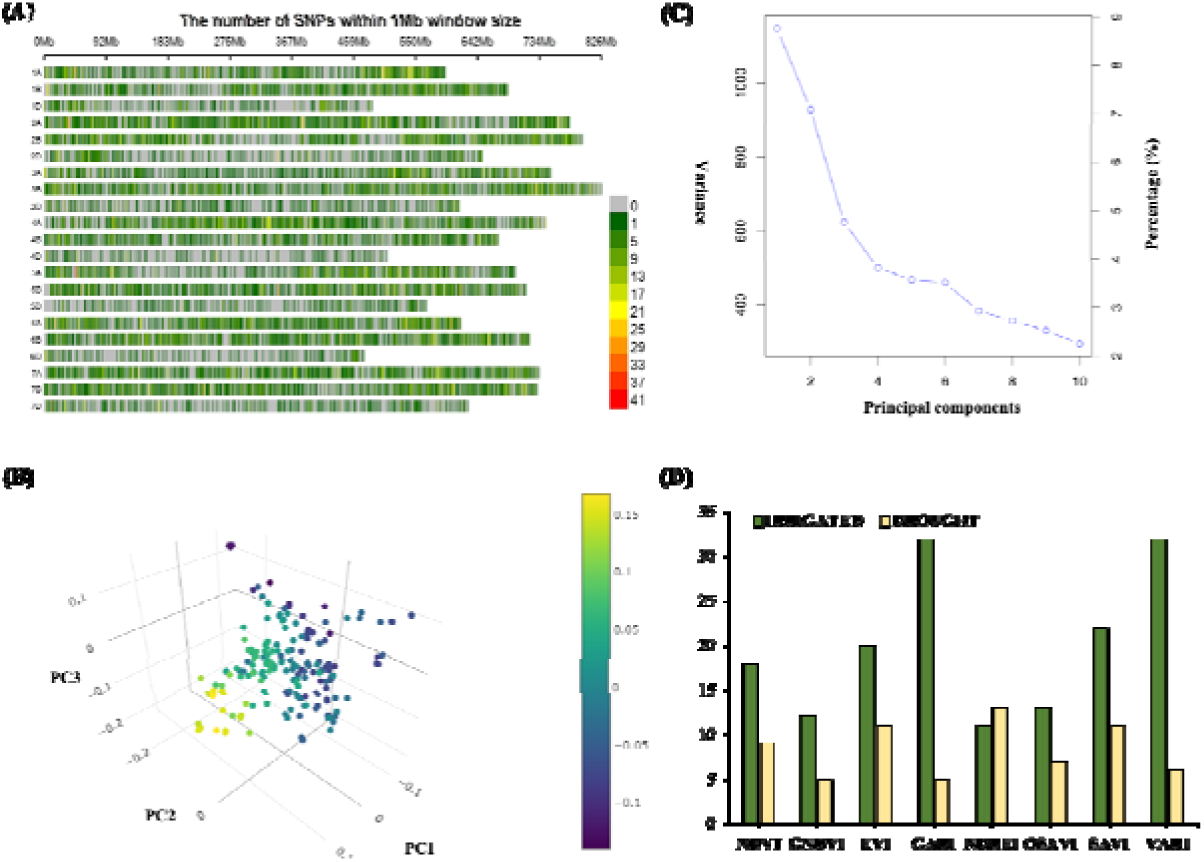
Population genomic analyses based on 23,897 high-quality SNP markers. A) SNP density on each chromosome, B) PCA plot to evaluate the population structure, C) eigen-values based graph for the selection of PCA, and D) number of significant SNPs under irrigated (green bars) and drought conditions (yellow highlighted bars).

**Figure 5:**
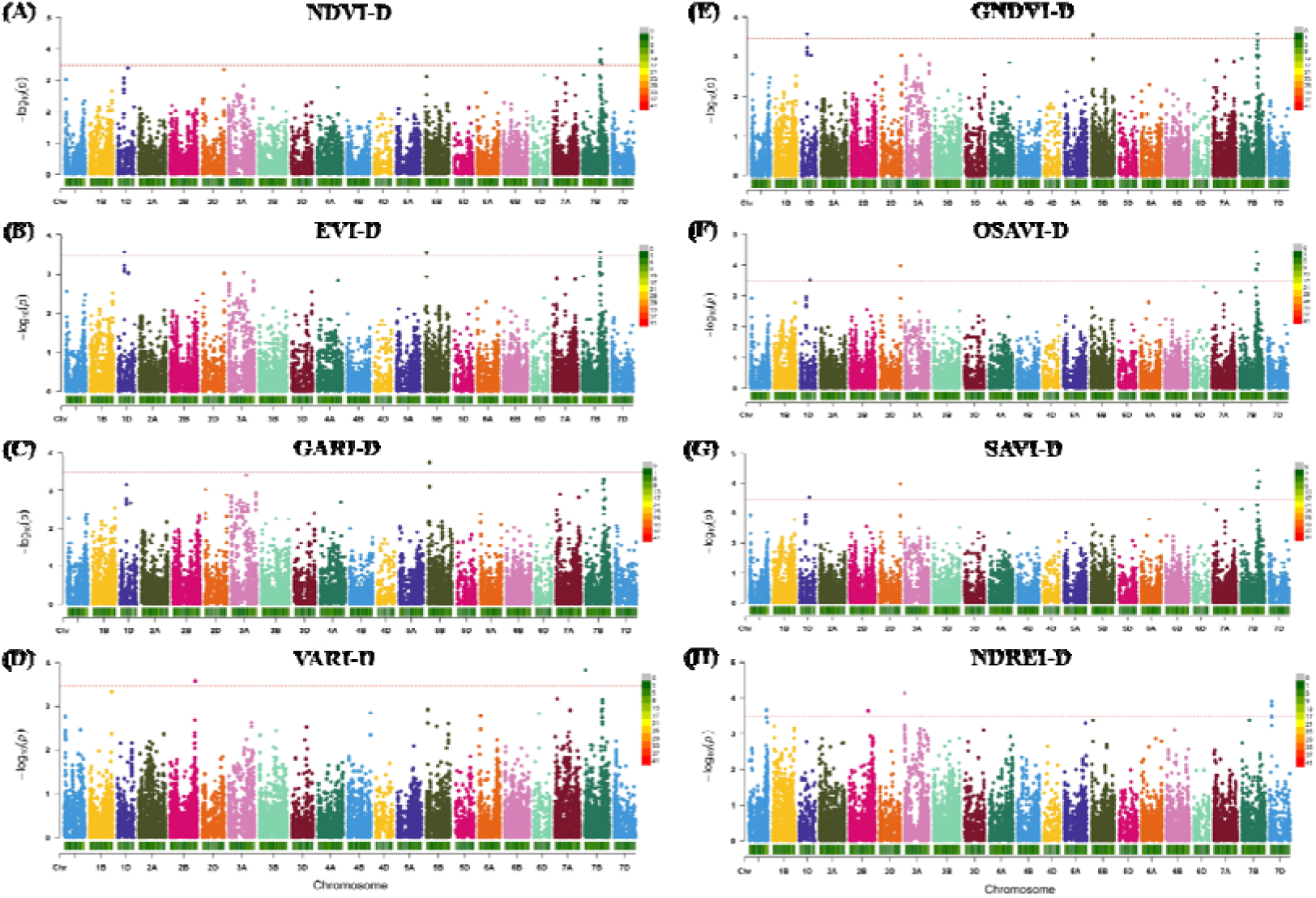
Marker-traits association for eight VIs under drought treatment. The haplotype analysis of pleotropic loci further identified several SNPs significantly associated with vegetation indices under drought stress with obvious statistical allelic effects (Fig 5). It helped us to understand that which allele is important or favourable allele at particular locus. Based on favourable allele information, we are able to estimate the favourable allele frequency (FAF) in each particular cultivars. Based on FAF, 26 lines were selected having more than 50% FAF (Table S24). FAF acquired the genome-based pre-breeding information, that can be important for drought breeding program. In future using these SNPs, breeders can develop the KASP markers, validate in their germplasm and utilize them for marker-assisted breeding.

**Figure 6:**
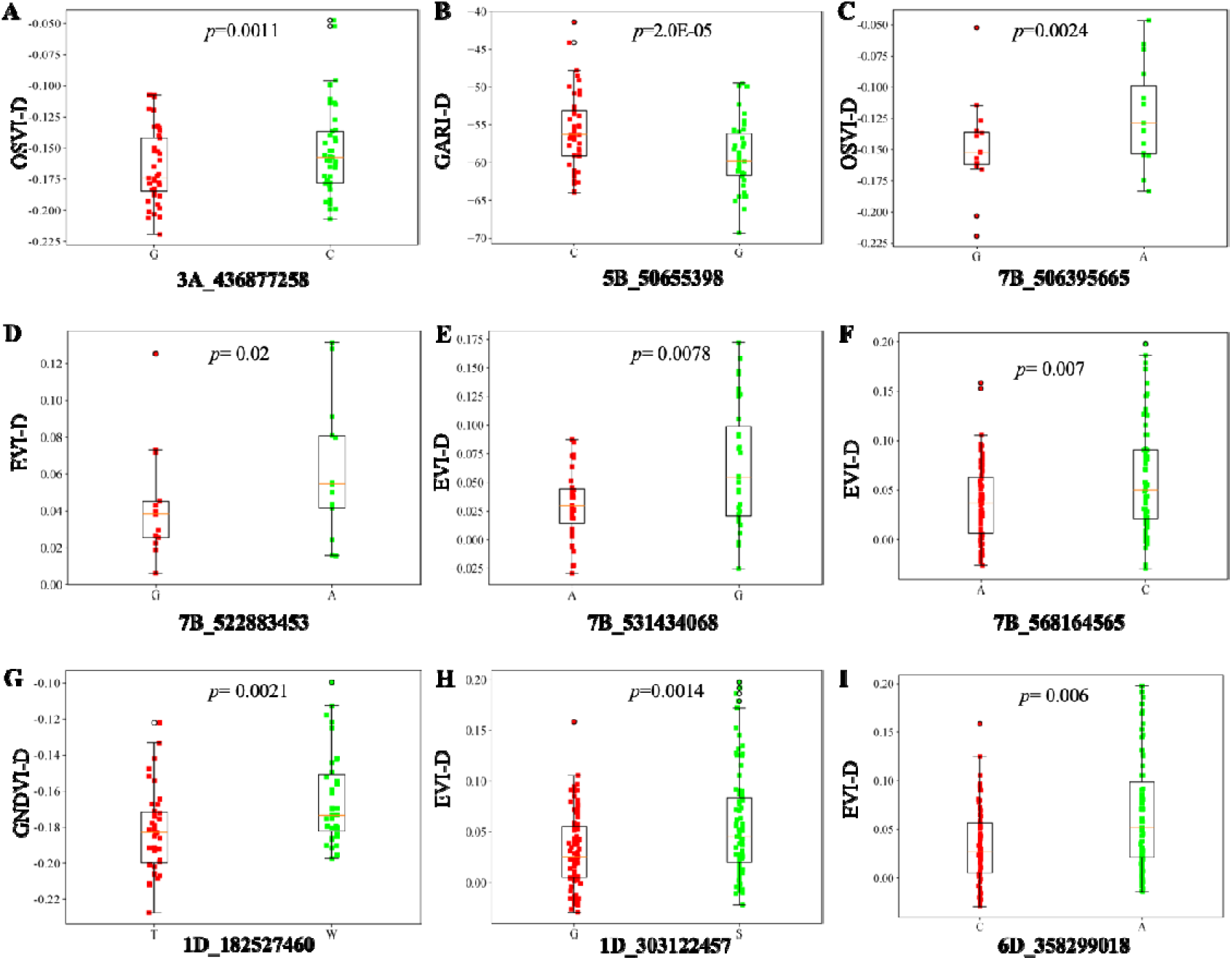
Haplotypes with significant allelic effect on VIs. Green colour box represent the favourable alleles have higher value for VIs under drought. Red highlights the unfavourable alleles with lower value. P-value represent the significance variation among both alleles for each trait.

### Population Genomics Analysis

Population genomics and GWAS analyses revealed substantial genetic variation across the wheat genome, with subgenome B harbouring the highest SNP density and subgenome D the lowest (Figure 4). Several SNP-rich hotspots were identified on chromosomes 2B, 3B, 5B, and 6B, while SNP-poor regions were prevalent in subgenome D.

PCA was carried out using 23,897 high-quality SNP markers. The first three components (PC1, PC2, and PC3) were visualized in a three-dimensional plot (Figure 4B), explaining 8.9%, 7.1%, and 4.8% of the total genetic variation, respectively. The distribution of wheat genotypes across this plot showed no distinct clustering pattern, indicating that the historical cultivars form a largely mixed population. Based on the variance explained, the first four PCs were incorporated as covariates in the GWAS model (Figure 4C), as a sharp decline in variance was observed after PC4.

### Identification Genetic Loci Linked with VIs

GWAS was carried out using the FarmCPU model to identify significant SNPs associated with VIs under both irrigated and drought conditions. The MLM model was employed to validate marker-trait associations. A total of 228 QTNs were identified across eight Vis under both environments, of which 67 were specifically detected under drought condition. A total of 26 pleiotropic loci were identified in this study, key genomic regions associated with multiple VIs. Among these, 10 loci detected under drought conditions were prioritized as promising targets for the genetic improvement of drought resilience in wheat.

A locus located at 18.25 Mb on chromosome 1D (1D_182527460) exhibited a positive effect on both GNDVI and OSAVI, indicating its potential role in maintaining canopy greenness and promoting biomass accumulation under drought. Similarly, SNPs 1D_303122457 (30.31 Mb) and 2D_645941939 (64.5 Mb) showed positive associations with EVI and NDVI. On chromosome 3A, the SNP 3A_436877258 (43.68 Mb) demonstrated a strong negative effect on GARI and OSAVI. Likewise, SNP 5B_50655398 (50.6 Mb) negatively influenced GARI and GNDVI. In contrast, SNP 6D_358299018 displayed a positive pleiotropic effect on both EVI and SAVI. Notably, chromosome 7B was identified as a pleiotropic hotspot with four significant SNPs (7B_506395665, 7B_522883453, 7B_531434068, and 7B_568164565) located at 50.63 Mb, 522.8 Mb, 53.14 Mb, and 56.81 Mb, respectively. The 7B_506395665 and 7B_522883453 positively regulated OSAVI, SAVI, EVI, and GARI. Remarkably, 7B_568164565 emerged as the most pleiotropic locus which influenced EVI under drought stress and NDVI, GNDVI, NDVI, and SAVI under irrigated conditions.

### Gene Annotation of Pleotropic Loci

Genes located closest to the significant SNPs were designated as possible putative candidate genes associated with vegetation indices under drought conditions. Annotation of 10 pleiotropic loci associated with VIs under drought identified 42 annotated genes with known functional relevance with drought tolerance (Table S25). Functional annotation of these genes examined two reported genes in wheat related to drought tolerance (Table S26). SNP 7B_522883453, has pleotropic effect for VI, GARI, GNDVI, NDVI, OSAVI, SAVI, NDVI, under drought stress was linked to *TraesCS7B01G287100.1* (*TaSnRK3.23B*) that is well characterized in drought tolerance. Pleiotropic locus, 7B_568164565, associated with NDVI and SAVI under drought stress, was functional annotated as *TraesCS7B01G318200.1* (*TaSAUR75*) known for its regulatory role in vegetative growth and stress responses. On chromosome 1D, the SNP 1D_303122457 (associated with NDVI, SAVI, EVI under drought) was linked to *TraesCS1D01G217600.1*, encoding an *Hsp70s* genes, which plays a protective role under abiotic stress (Table S26).

While among the remaining 7 pleotropic loci, 6 pleotropic loci were annotated with ortholog genes that have important role under drought stress (Table S26). These ortholog genes belong to different crops like *Arabidopsis*, rice, maize and soybean. Another SNP on 1D (1D_182527460), associated with OSAVI, GNDVI under drought, was annotated with *TraesCS1D01G136900.1*, a Ras-related protein (*AtRabE1c).* On chromosome 2D, SNP 2D_645941939 (NDVI-D, SAVI-D, EVI-D) was linked to *TraesCS2D01G589300.1*, which encodes *AtGSTU17* gene. The locus 3A_436877258, associated with OSAVI-D, GARI-D, was annotated with *TraesCS3A01G233800.1*, *OsTPKb* related to Potassium channel gene function. The SNP 5B_50655398 (OSAVI-D, GARI-D, GNDVI-D, NDREI-D) corresponded to *TraesCS5B01G045300.2*, Zinc finger CCCH domain-containing protein 4 (*GmZF351*), likely involved in stress-responsive gene expression. On chromosome 6D, SNP 6D_358299018 (EVI-D, SAVI-D) was annotated with *TraesCS6D01G254000.1*, monodehydroascorbate reductase encoding *ZmMDAR1* gene. SNP 7B_531434068 (GARI-D, GNDVI-D, NDVI-D, OSAVI-D, SAVI-D) was associated with TraesCS7B01G293900.1, BZIP transcription factor ortholog of *OsbZIP62* gene (Table S26).

Annotated genes were further diagnosed for publicly available transcriptome data (http://barleyexp.com/exp.php). Transcriptome profiling revealed that among the 42 annotated genes only six genes *i.e*., *TraesCS7B01G287100.1, TraesCS7B01G318200.1, TraesCS1D01G217600.1*, *TraesCS1D01G136900.1*, *TraesCS3A01G233800.1,*

*TraesCS6D01G254000.1* might be putative candidate genes (Table 1 and Figure 7). These possible putative candidate genes were further validated by quantitative PCR (qPCR) either these genes also showed differential gene expression among drought tolerant and susceptible cultivars. Drought tolerant *i.e.,* Parwaz-94 and Gulzar-19 and susceptible *i.e.,* AARI-10 and Wafaq-23 genotypes were selected based on favourable allele frequency. For this purpose two genes *i.e., TraesCS1D01G217600.1* and *TraesCS6D01G254000.1* with higher transcriptome value were selected for qPCR. Relative expression of these genes revealed differential expression profiling between tolerant and susceptible cultivars. Side by side both genes showed upregulation under drought stress (Figure 7B and 7C). These results suggested that these two genes are candidate genes and have their role in drought tolerance mechanism. In future characterization of these genes in wheat will help to understand the genetic mechanism and their role in drought tolerance.

**Figure 7:**
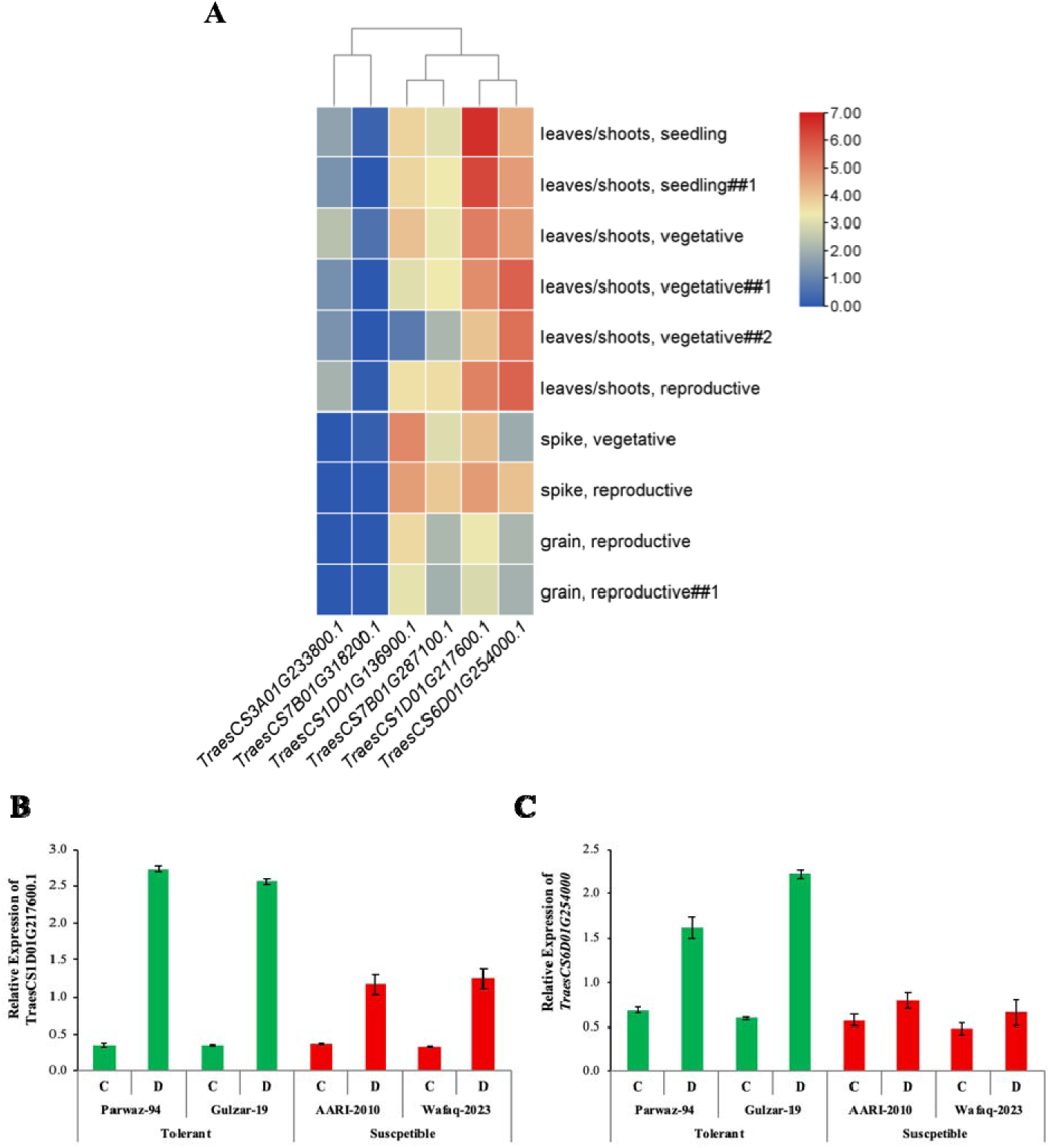
Identification of candidate genes using expression profiling approach. **A)** Transcriptome profiling of six putative candidate genes at different plant tissues and development stages based on tpm value. **B)** Relative expression of *TraesCS1D01G217600.1* in drought tolerant highlighted with green bars and susceptible cultivars demonstrated with red bars under normal irrigation (C) and water-limited condition (D). **C)** Relative expression of *TraesCS6D01G254000.1* in drought tolerant highlighted with green bars and susceptible cultivars demonstrated with red bars under normal irrigation (C) and water-limited condition (D).

**Table 1:** Candidate gene with respective pleotropic loci.

| S. No. | SNP | CHROM | POS | Traits | Transcript ID | Gene physical position | Ortholog Gene | Annotated genes |
| --- | --- | --- | --- | --- | --- | --- | --- | --- |
| 1 | 1D_182527460 | 1D | 182527460 | OSAVI-D, GNDVI-D | TraesCS1D01G136900.1 | 184470873 | *AtRabE1c* | Ras-related protein |
| 2 | 1D_303122457 | 1D | 303122457 | NDVI-D, SAVI-D, EVI-D | TraesCS1D01G217600.1 |  | *HSP70* | Chaperonin 10 Kd subunit |
| 3 | 3A_436877258 | 3A | 436877258 | OSAVI-D, GARI-D | TraesCS3A01G233800.1 | 436466697 | *OsTPKb* | Potassium channel |
| 4 | 6D_358299018 | 6D | 358299018 | SAVI-D, EVI-D | TraesCS6D01G254000.1 | 358225193 | *ZmMDAR1* | Monodehydroascorbate reductase |
| 5 | 7B_522883453 | 7B | 522883453 | EVI-D, GARI-D, GNDVI-D, NDVI-D, OSAVI-D, SAVI-D, NDVI-D | TraesCS7B01G287100.1 | 522719981 | *TaSnRK3.23B* | Serine/threonine-protein kinase |
| 6 | 7B_568164565 | 7B | 568164565 | NDVI-D, SAVI-D | TraesCS7B01G318200.1 | 568106785 | *TaSAUR75* | SAUR-like auxin-responsive family protein |

## Discussion

Plants express a remarkable diversity of traits, ranging from structural and physiological features to developmental patterns, that together form highly complex phenotypes (Kaya, 2025, Wright et al., 2022). Capturing this complexity has long been a challenge, but recent developments in image-based phenotyping have opened new possibilities. By enabling automated extraction of features from digital images, these technologies are helping overcome one of the main constraints in plant phenotyping: the slow, labor-intensive nature of conventional measurements (Atefi et al., 2021). The growing range of HTP platforms now makes it possible to follow plant growth and development in fine detail and across different developmental stages (Li et al., 2021, Khuimphukhieo and da Silva, 2025, Raza et al., 2025).

With the rapid expansion of phenomics, there is an increasing need for analytical approaches that can cope with large, complex image datasets, extract meaningful traits, and integrate these with other omics layers to provide deeper biological insights (Yang et al., 2021). In this study, we applied such an approach by combining image-derived phenotypic data with GWAS. From the UAV imagery, we derived eight VIs by analyzing canopy-reflected light across multiple wavelengths. Measurements were taken at early growth stages under both irrigated and drought conditions. This approach ensured pixel consistency during VI calculation, allowing us to detect early drought responses and assess genotype-by-environment (G×E) interactions. Compared with traditional methods, this strategy provided a more accurate and efficient way to predict complex physiological traits.

As expected, drought stress had a marked impact on chlorophyll content, photosynthetic performance, and canopy structure, which in turn influenced all VIs. NDVI, a widely used measure of chlorophyll and canopy density (Yan et al., 2025), dropped sharply under drought, approaching zero at both TP1_d and TP2_d, while remaining relatively stable (0.2–0.5) under irrigation. Similar patterns have been reported by (Yan et al., 2025). GNDVI, which is more sensitive to chlorophyll variation than NDVI (Yan et al., 2025), showed negative values (−0.2 to 0.1) in drought-stressed plants, indicating substantial pigment loss, while control plants maintained positive values (∼0.1). These findings align with (Hassan et al., 2019), the latter highlighting GNDVI’s advantage for early drought detection. Other indices reflected related changes.(Matsushita et al., 2007), was slightly lower in droughted plants, suggesting reductions in leaf area and biomass. GARI, used to minimize atmospheric interference in stress detection (NilJu et al., 2025). Indices designed to correct for soil background, such as SAVI (Kareem et al., 2023) and OSAVI (Rondeaux et al., 1996), also decreased significantly under drought, indicating reduced vegetative vigour, consistent with the results of (Pipatsitee et al., 2023). VARI, which measures greenness in the visible spectrum (Qubaa et al., 2021), fell notably in drought-stressed plants, echoing findings by (Rockstad et al., 2024) on drought-induced leaf senescence in wheat. Our results show that UAV-based multispectral indices can provide sensitive, early-stage indicators of drought-induced changes in wheat.

In this study, VIs displayed a wide range of variation along with consistently high heritability. Such traits are particularly valuable for improving the power of GWAS to detect meaningful genetic associations. Previous work has shown that when target traits exhibit both large variation and high heritability, the likelihood of identifying significant loci is greatly improved. Our marker–trait analysis revealed 26 pleiotropic loci distributed across chromosomes under both control and drought conditions. Of these, 10 loci showed strong and specific associations with VIs across chromosomes 1D, 2D, 3A, 5B, 6D and 7B under drought stress. (Rabbi et al., 2021) reported a novel QTL on chromosome 2D linked to drought tolerance, and other studies have identified NDVI-associated loci on 2D under moisture stress. (Hassan et al., 2021) reported several vegetation index–related QTLs on chromosome 3A using UAV-based multispectral imaging, highlighting its role in controlling senescence and canopy greenness. (Vukasovic et al., 2024) reported a functional locus on chromosome 5B associated with root biomass, transpiration, and nitrogen uptake, all of which influence vegetation indices under drought. (Puttamadanayaka et al., 2020) reported significant NDVI-related QTLs on chromosome 6D in bread wheat under water-deficit conditions, suggesting its involvement in maintaining canopy greenness and yield components during stress. (Condorelli et al., 2018) reported major QTL hotspots related to NDVI and stay-green trait on chromosome 7B under drought conditions. This finding points to the presence of drought-responsive genetic regions that could be critical in breeding wheat varieties with enhanced climate resilience. Importantly, the loci identified here might have strong potential for practical breeding. By converting them into KASP markers, breeders could incorporate them into marker-assisted selection pipelines (Liu et al., 2023, Rubab et al., 2023). Such markers are especially useful for trait pyramiding, allowing favorable alleles from different loci to be combined efficiently. This would support the development of wheat cultivars that not only yield well but also maintain performance under challenging environmental conditions.

The identification of putative candidate genes near significant SNPs provides valuable insights into the genetic basis of VIs under drought stress in wheat. The SNP 1D_182527460, which was significantly associated with OSAVI and GNDVI under drought stress, was annotated with the *AtRabE1c* gene, a member of the Rab GTPase family known to play critical roles in vesicle trafficking, stress signalling, and plant adaptive responses. (Chen et al., 2021) reported that *TaRabGTPases*, including *TaRabE1c* homologs, are involved in enhancing drought tolerance by regulating cellular membrane trafficking and improving stress-related physiological efficiency in wheat The *Hsp70* gene, associated with SNP 1D_303122457 functions as a molecular chaperone, protecting proteins from denaturation under stress. The co-location of an *Hsp70* with SNPs affecting NDVI-D, SAVI-D, EVI-D is consistent with Hsp70-mediated maintenance of photosynthetic apparatus stability and pigment-related changes that alter vegetation indices under stress (Yu et al., 2015). In wheat, it has been linked to improved drought and heat tolerance (Kou et al., 2023). The link to EVI and NDVI under drought treatment in our analysis indicates the common genetic regulation for stress signalling in canopy under drought stress. The glutathione-S-transferase (*AtGSTU17* orthologue) was located near SNP 2D_645941939 is another possible putative candidate gene. Functional characterization of glutathione-S-transferase gene in *Arabidopsis, Populus, grapevine* showed their role in drought resilience (Chen et al., 2012; Nerva et al., 2021; (Niu et al., 2024). Glutathione homeostasis, ABA signalling and stomatal/ROS regulation; mutation or altered expression of *GSTU17* changes plant water-loss traits and oxidative status under drought and salt stress (Chen et al., 2012). The locus 3A_436877258, associated with GARI and OSAVI, annotated to a vacuolar two-pore KlJ channel (*OsTPKb* orthologue). Overexpression and functional characterizations of *TPK/TPKb* channels in rice indicate these channels contribute to vacuolar KlJ homeostasis and osmotic balance, processes central to cell turgor maintenance during drought (Ahmad et al., 2016). Variation at a *TPK* locus could therefore alter leaf turgor and pigment properties that influence GARI/OSAVI signals. The CCCH-type zinc-finger (*GmZF351-like*) was located near 5B_50655398. Recent functional work in soybean demonstrates that *GmZF351* is stress-inducible and that overexpression improves salt and drought tolerance, likely through transcriptional reprogramming of downstream stress-responsive genes (Wei et al., 2023). A zinc-finger at this locus therefore provides a regulatory explanation for co-variation of GARI, OSAVI, NDREI and GNDVI under drought stress.

The 6D locus (6D_358299018) annotated to MDAR family members (*ZmMDAR1-like*) involved in the ascorbate–glutathione cycle. MDAR enzymes recycle monodehydroascorbate to ascorbate and are integral to the cellular antioxidant network (Shin et al., 2013). Previous studies show MDAR activity and expression respond to drought/heat and modulate oxidative damage (Liu et al., 2012). The variations in MDAR locus associated with VIs indicates sensitive to chloroplast redox state and pigment degradation under stress. Pleiotropic loci on 7B, 7B_506395665, 7B_522883453, and 7B_531434068, point to regulators of cell-wall modification (*TBL*), *SnRK3* kinase family members (*TaSnRK3.23B*), and a cluster including a *bZIP* transcription factor (*OsbZIP62-like*) and expansin/pollen-allergen family proteins, respectively. TBL proteins modify xylan acetylation and cell-wall properties that influence leaf optical properties (Gao et al., 2017), while *SnRK3* kinases and *bZIP* TFs are central to ABA and osmotic stress signalling (Takahashi et al., 2020, Banerjee and Roychoudhury, 2017). *OsbZIP62* and *TaSnRK3* family members modulate ABA-responsive gene networks and oxidative tolerance; their co-localization with VIs associated SNPs indicates genotype-dependent modulation of ABA signalling and cell-wall structure contributes to observed greenness of plants under drought. The pleiotropic locus 7B_568164565 linked with NDVI and SAVI, annotated as *TaSAUR75*. Functional overexpression studies of *TaSAUR75* indicate reduced ROS accumulation and enhanced tolerance to drought and salt when expressed in heterologous systems (Lv et al., 2022). The presence of *TaSAUR75* at a locus suggests that allelic variation in *SAURs* influences physiological status in ways that are captured by multispectral indices. These co-locations connect variation in VIs to canonical stress pathways including molecular chaperoning (*Hsp70*), redox and antioxidant recycling (*GST, MDAR*), ion/osmotic homeostasis (*TPK*), transcriptional reprogramming (zinc-finger, *bZIP, SnRK*), cell-wall modification (*TBL*) and growth regulation (*SAUR*).

Transcriptome profiling of these genes help us to select the most possible putative candidate gene. Based on transcriptome profiling six genes i.e., *TraesCS7B01G287100.1 (TaSnRK3.23B), TraesCS7B01G318200.1 (TaSAUR75), TraesCS1D01G217600.1 (HSP70s)*, *TraesCS1D01G136900.1 (AtRabE1c)*, *TraesCS3A01G233800.1 (OsTPKb),* and *TraesCS6D01G254000.1 (ZmMDAR1)* showed differential gene expression which suggests that might be these genes play important role in drought tolerance. Further, qPCR analysis of two genes *TraesCS1D01G217600.1 (HSP70s)* and *TraesCS6D01G254000.1(ZmMDAR1)* reveals that these genes have their role in drought tolerance. In future the functional validation of these genes will help to understand the genetic mechanism underlying these candidate genes. This study provides targets for breeding drought-resilient germplasm that maintain favourable spectral signatures of canopy.

## Declarations Clinical trial

Not applicable.

## Acknowledgments

The authors are thankful to Pakistan’s Research Institutes for providing Pakistan’s historical cultivars. The authors extend their appreciation to the National Institute for Genomics and Advanced Biotechnology (NIGAB), NARC and Chengdu Institute of Biology (CIB), CAS for joint research venture.

## Author Contributions

Conceptualization, Muhammad Kashif Naeem, and Muhammad Ramzan Khan; Data curation, Muhammad Asad Ullah, Fara Muqadas; Formal analysis, Muhammad Asad Ullah, Fara Muqadas, Atiq ur Rehman; Funding acquisition, Muhammad Ramzan Khan, Jing Chen; Investigation, Zahid Mahmood, Muhammad Sajjad, Muhammad Arshad Javed; Methodology, Muhammad Kashif Naeem, Muhammad Sajjad, Shoaib ur Rehman and Javaria Tabassum; Supervision, Muhammad Ramzan Khan, Muhammad Arshad Javed; Validation, Jing Chen, Muhammad Ramzan Khan; Visualization, Muhammad Kashif Naeem, Muhammad Asad Ullah and Shoaib ur Rehman; Writing – original draft, Muhammad Asad Ullah, Fara Muqadas and Muhammad Kashif Naeem; Writing – review & editing, Zahid Mahmood, Jing Chen, and Muhammad Ramzan Khan. All authors have read and agreed to the published version of the manuscript.

## Funding

The authors extend their appreciation to the Researchers Supporting Project Sichuan Science and Technology Support Program, China (2025HJRC0044), the National Key R&D Program of China (2024YFD1201203). The research was also funded by the Public Sector Development Program (PSDP) “Sino-Pak Agricultural Breeding Innovations Project for Rapid Yield Enhancement” (PSDP-857) at the National Institute for Genomics and Advanced Biotechnology (NIGAB), NARC.

## Data Availability Declaration

The data that supports the findings of this study are available in the supplementary material of this article.

## Ethical approval

Not applicable.

## Conflict of Interest

The authors declare that the research was conducted in the absence of any commercial or financial relationships that could be construed as a potential conflict of interest.

## Notes

### Competing Interest Statement

The authors have declared no competing interest.

